# Helical Growth of Twining Common Bean is Associated with Longitudinal, Not Skewed, Microtubule Patterning

**DOI:** 10.1101/2025.05.27.656463

**Authors:** Angelique A. Acevedo, Mariane S. Sousa-Baena, Joyce G. Onyenedum

## Abstract

Organ chirality in plants has been linked to cytoskeletal organization, as demonstrated in *Arabidopsis thaliana* twisted mutants, where left-skewed cortical microtubules are associated with right-handed twisting, and vice versa. While this phenotype seemingly mirrors vining activity, this hypothesis remains understudied within naturally twining plants. Qualitative observations identified skewed microtubules in the twining stem of *Ipomoea nil* vine, suggesting parallels with *Arabidopsis* studies. To further investigate organ chirality in twining plants, we used common bean vine (*Phaseolus vulgaris* L.) to examine the relationship between microtubule orientation, cell morphogenesis, and the right-handed twining phenotype via immunolabeling techniques. Here, we report a transition from mixed microtubules orientations in emergent and elongating internodes to a predominance of longitudinal microtubules in straight and twined stem segments post-elongation. Additionally, we report a distinction in epidermal cell shapes, where the straight portions of the stem consist of lobes with rectangular cells and furrows comprised of round cells, while the twined portions are comprised of cells that are relatively more rectangular and stretched. We propose that these orientations reflect dynamic microtubule responses to external stimuli and growth cues, such as tensile stresses from climbing or tissue expansion. Taken together, these findings highlight dissimilarities between twisting *Arabidopsis* mutants and naturally twining plants.

**Highlight:** Longitudinally arranged cortical microtubules found along the coiled stems of common bean vine highlight disparities between the directional growth of well-documented twisted *Arabidopsis* mutants and naturally twining plants.

## Introduction

Twining vines provide an opportunity to understand the developmental mechanisms underlying rapid directional growth in plants, given the uniquely dynamic movements characterizing their growth form (Sousa-Baena et al., 2021). While it is known that most twining vines coil in a right-handed fashion (Edwards et al., 2007; Zhou et al., 2019), the cellular underpinnings that constitute handedness in naturally twining species remain understudied. At present, insights into organ handedness have primarily come from microtubule mutants in the model species, *Arabidopsis thaliana* (L.) Heynh. These studies’ converge on a growing consensus that twisted organ growth (twisting roots, hypocotyl, leaf petioles, and petals) is correlated with skewed microtubules in the following way: left-handed microtubules serve as the track for left-handed cellulose microfibrils, which permits cells to grow in a right-handed direction, and vice versa. This pattern has been found in several mutants, including *SPIRAL1 (spr1)*, *tortifolia1/SPIRAL2 (tor1/spr2)*, *tortifolia2 (tor2)*, and *lefty* mutants (Furutani et al., 2000; Thitamadee et al., 2002, Buschmann et al., 2004, Nakajima et al., 2004, Shoji et al., 2004, Buschmann et al., 2009). However, exceptions to this pattern exist, such as the *SKU6/SPIRAL1 (spr1-6/spr1-1)* mutant, where right-handed twisting organs exhibited transverse and right-handed microtubules (Sedbrook et al., 2004). More generally, skewed microtubules have also been observed in the straight elongation zone of roots (Liang et al., 1996). These exceptions emphasize the complex relationship between microtubules and organ-level twisting as a result of typical microtubule variability and dynamics.

While other components of the cell wall, like rhamnogalacturonan (RG-I) pectins are shown to affect plant cell patterning, as shown in the *RHM1 (rmh1-1) Arabidopsis* mutant (Saffer et al., 2017), most research has focused on the link between cellulose microfibrils and cell expansion, because cellulose is the primary load-bearing element of the cell wall, constraining maximal cell expansion perpendicular to their net orientation (Baskin, 2005; Anderson et al., 2010, Buschmann and Borchers, 2020). However, this net constraint is often transformed by the dynamic behavior of cortical microtubules, which self-organize beneath the plasma membrane to guide cellulose synthase complexes and orient the deposition of cellulose layers in the plant cell wall (Heath, 1974; Paredez et al., 2006; Duncombe et al., 2022). When microtubule orientation—and consequently cellulose arrangement—is randomized across the cell axis, cell expansion is isotropic (uniform), whereas unidirectional microtubule arrangement is termed anisotropic (directional) due to unequal yielding of the cell wall (Green, 1962). Generally, plant cells can undergo both isotropic and anisotropic growth. Prolonged isotropic expansion typically produces rounded cells, and anisotropic expansion produces elongated cell morphologies, yet it is the combination and assortment of these uniquely shaped cells forming plant organs that makes them an area of interest for investigating twisted organs.

Despite the well-documented role of microtubule arrangement in modulating cell anisotropy and organ-twisting of *Arabidopsis* mutants, its role in naturally twining species remains understudied. Sousa-Baena et al. (2021) provided an initial qualitative analyses performed on a twining vine, *Ipomoea nil* (L.) Roth, which revealed skewed microtubule arrangement in fully twined internodes, drawing parallels to the work completed in *Arabidopsis* studies. In this study, we aim to clarify and deepen our understanding of the relationship between cortical microtubules and right-handed twining by using common bean vine, *Phaseolus vulgaris* L. (Fabaceae). We report the counterintuitive finding that a longitudinal––rather than a left-or right-skewed––microtubule arrangement dominates in twined internodes. Furthermore, we identified a link between stem segment morphology and cell shape, whereby straight stem segments display alternating lobes with elongated rectangular cells and furrows with rounded cells, and twined internodes consisted of cells possessing a stretched quality. Our results begin to highlight an important distinction between the development of twining in vines, such as when helical growth observed in common bean requires internodes to elongate and grow in contact with a support, versus twisted *Arabidopsis* mutants, where twisting occurs at the level of the cell.

## Methods

### Plant Materials and Growing Conditions

In this study, we selected *Phaseolus vulgaris* L. RIL L88-57 as a model system to investigate the cellular mechanisms underlying twining. To improve the success and uniformity of seed germination, all seeds were imbibed for 24 hrs in 20% (w/v) of PEG 4000 in the dark and thoroughly rinsed with deionized water. After rinsing, seeds were transferred to Petri dishes containing dampened 90 mm diameter Whatman™ filter paper (GE Healthcare Life Sciences, Cat No. 1001-090) for 2-4 days at room temperature in indirect bright light until the radicle emerged.

Germinated seeds were then transferred to 475 mL (4-inch) pots containing Lambert LM-111 All Purpose soil mix (Cat No. 664980 2510). L88-57 plants were grown under 12-hour light conditions at ∼130 μ*mol*/*m*”/*s* PAR, followed by a 30-minute ramp-down period to a 12-hour dark cycle and repeating with a 30-minute ramp-up back to the light cycle. Pots were positioned ∼76 cm below LED light panels in Environmental Growth Chambers (EGC models GR48 and SLR-90) with conditions set to 50% relative humidity and a temperature of 24**°** C. Plants were also supplemented with 75 ppm of Jack’s Professional 20-20-20 General Purpose fertilizer containing micronutrients.

### Selection of Morphological Stages

To understand the relationship between microtubule angles, cell morphogenesis, and plant morphology, we sampled plants at key biomechanical stages (sensu Onyenedum et al., 2025): Stage 1 (S1) plants are erect seedlings with embryonic leaves; Stage 2 (S2) plants have a slight bend of the epicotyl with internode development; Stage 3 (S3) is characterized by the small revolutions of the shoot apex, known as ‘regular circumnutation’; Stage 4 (S4) marks a period of internode elongation and ‘exaggerated circumnutation’, and stage 5 (S5) is defined by a plant’s twined position. We focused on stages 2, 3, and 5 for their distinct biomechanical features and habits, representing critical developmental periods necessary for twining.

At these stages, we selected three internodes to investigate, in addition to the hypocotyl, which served as a comparable reference to previous microtubule studies conducted on *Arabidopsis* hypocotyls. Selected internodes provided unique insights into the basal, central, and twined regions of a developing plant. Internode 1 remained stationary and relatively short throughout stages 2, 3, and 5. Internode 3 acted as a transition zone, facilitating circumnutational movements observed in stage 3 before becoming straight and stationary at stage 5. Similarly, internode 6 participated in circumnutation at stage 3, but settled into a twined stationary position at stage 5.

### Microtubule Immunolabeling

To characterize microtubule (MT) patterning across key biomechanical stages of twining, we labeled epidermal peels with an anti-α-tubulin antibody and tracked MT arrangement via confocal microscopy. Epidermal peels were prepared by nicking the surface of a fresh internode with an Astra Double Edge razor, then gently grasping the epidermal flap with blunt-ended tweezers and pulling straight downwards as described in Median et al., (2022). A minimum number of three individuals were sampled, focusing on their hypocotyl and internodes 1, 3, and 6 at three key developmental stages: stage 2 (S2), stage 3 (S3), and stage 5 (S5).

Microtubules were labeled using protocols adapted from Wasteney et al. (1997) and Cytoskeleton: Methods and Protocols (Celler et al., 2016, p 155-184) with modifications for *Phaseolus vulgaris* L. tissue. Thin epidermal peels from selected internodes were fixed for 40 minutes in a 0.5% glutaraldehyde and 1.5% formaldehyde solution before being washed three times for 10 minutes in PMET buffer (50 mM PIPES, 5mM EGTA, 1mM magnesium sulfate, and .05% Triton X-100, pH 7.2). The samples were then placed between two plain non-frosted microscope slides (VWR, Cat No. 48300-026), secured by two heavy-duty spring clamps, and flash-frozen by being dipped in a bath of liquid nitrogen with tongs. Once frozen, the slides were directly transferred to a – 80°C aluminum block and crushed with enough force to generate noticeable cracks along the peel. The slides were immediately pulled apart and placed onto a room-temperature aluminum block to thaw. This freeze-thaw cycle was repeated by dipping the slides into liquid nitrogen a second time. Samples were then incubated in a cell wall digestion enzyme solution for 30 minutes (0.05% pectolyase from *Aspergillus japonicus*, 0.4M D-mannitol, 1% BSA in PBS). Following the digestion solution, samples were washed three times for 10 minutes in PMET buffer and left to incubate for 3 hours at room temperature in permeabilization buffer on a variable-speed rocker (1% Triton X-100 in PBS, pH 7.5).

Furthermore, samples were washed three times for 10 minutes in PBS solution and transferred to a sodium borohydride solution for 20 minutes (1 mg/mL sodium borohydride in PBS). The sodium borohydride solution was discarded and replaced with a blocking buffer for 30 minutes (1% BSA, 50 mM glycine in PBS). A primary antibody (AB) solution of mouse anti-α-tubulin B 512 (Krackeler Scientific Inc. (Albany, NY), Cat No. 45-T6074-200UL-EA) in blocking buffer was used at a 1:1000 dilution to incubate peels for ∼14-16 hours in 1.5 mL microfuge tubes at 4°C. Subsequently, samples were rinsed five times for 10 minutes each with an incubation buffer in petri dishes on a variable-speed rocker (50 mM glycine in PBS). They were then placed into a blocking buffer for 30 minutes. Next, a secondary antibody (AB) solution, using Alexa 488-conjugated goat anti-mouse IgG (Thermo Fisher Scientific (Suwanne, GA), Cat No. A28175), was prepared at a 1:100 dilution in blocking buffer and used to incubate the samples at room temperature for 3 hours in the dark. Samples were then removed from the secondary AB solution and washed three times for 10 minutes in PBS on the rocker. Finally, peels were mounted in Citifluor™ AFI Antifading Mountant Solution with a coverslip and sealed the edges with fast-drying Sally Hansen top-coat nail polish.

Microtubule images were acquired using a Leica Stellaris 5/DMi8 confocal model with an inverted microscope at 20x and 63x/1.4 NA oil immersion objectives with a 2X zoom factor (excitation 488nm, emission 510 – 530nm range). Z-stacks were created to view cells from cortical to epidermal layers with a median noise reduction filter and later used to generate projections of the outermost periclinal side of the cell’s surface. These projections were imported into Adobe Photoshop and rotated to align the cells vertically, providing an upright longitudinal view. Next, the projections were processed with ImageJ using the FibrilTool plugin following Boudaoud et al. (2014)’s installation and fibril quantification process (https://doi.org/10.1038/nprot.2014.024). For each internode and individual (n≥3) across developmental stages, 30 epidermal cells were measured. Microtubules angled between 0°-22.5° and 157.5° – 180° degrees were labeled transverse, while angles 90° within a ±22.5° degree range were labeled longitudinal. Microtubules angled between 22.5° and 67.5° were categorized as right-skewed, and those between 112.5° and 157.5° were left-skewed. Binning in reference to Renou et al. (2024).

Barplot visualization of this data was completed using ggplot2, ggpubr, and ggpatterns packages (v3.5.1; Wickham, 2016; v0.6.0; FC and Davis, 2022; v1.1.4; Kassambara, 2023). Supplemental relative density curves were made using the ggridges package in RStudio (v0.5.6; Wilke, 2024).

### Propidium Iodide Staining and Classifying Cell Morphologies

To characterize cell morphogenesis through key biomechanical stages, we used propidium iodide (PI) to stain epidermal cells and morphologically segment their cell walls via ImageJ’s MorpholibJ plugin. PI was prepared as a stock solution of 1 mg/mL solution in deionized water (stored at 4°C) before being further diluted into a 10µg/mL working solution. Epidermal peels were made from fresh tissue and incubated in a 1.5 mL tube containing PI working solution for 30 minutes in the dark and on ice before being thoroughly washed three times for 2 minutes each. The tube was inverted 2 – 3 times and shaken using a Vortex-Genie 2.

Panoramic confocal images were generated by Leica Software using Leica Stellaris 5/DMi8 confocal model with a 20X dry objective at a 2X zoom factor. PI was detected at an excitation of 532nm and captured in the 590 – 630nm emissions range. The resulting images were imported and processed via ImageJ and the MorpholibJ plugin (Legland et al., 2016). This plugin was used to morphologically segment cells (with user adjustments using the ‘merge’ function) and label various ROIs with corresponding cell measurements: cell circularity, geodesic diameter, and cell area. Stomatal and trichome precursor cells were removed from the analysis. Once measurements were assigned to their corresponding ROI labels, the label edition feature was used to generate a spectral color map. Color maps reflected cell measurements like cell ‘area’ and were used to create a heat map with the LUT “Fire/Inferno” gradient.

Measurements were retrieved from ImageJ and converted into bar plots and boxplots using RStudio packages (ggplot2) and (RColorBrewer) to compare the distribution of cells in furrowed and lobed portions of internodes throughout development (v3.5.1; Wickham, 2016; v1.1-3; Neuwirth, 2022).

## Results

### Stem morphology shifts from base to shoot tip

To evaluate the relationship between cortical microtubules (CMTs) and the development of the right-handed twining phenotype, we first identified distinct morphological stages of common bean development *sensu* Onyenedum et al. (2025). From this study, stage 2 was selected to sample newly emergent seedlings developing its first internode (Fig. 1A, “Stage 2”); stage 3 to capture plants undergoing regular circumnutation of the shoot tip (Fig. 1A, “Stage 3”); and stage 5 to evaluate mature right-handed twining plants (Fig. 1A, “Stage 5”, 1B). We used epidermal markers such as stomata and trichomes to identify the periclinal side of peeled epidermal tissue (Fig. 2A, 2B). CMT arrays were categorized into one of four major orientation groups— transverse (0 – 22.5° and 157.5 – 180°), right-skewed (22.5 – 67.5°), longitudinal (67.5 – 112.5°), and left-skewed (112.5 – 157.5°) (Fig. 3A–2D). Internode elongation measurements are in Fig. S1 and Chi-square results are located in Table 1.

**Fig. 1:**
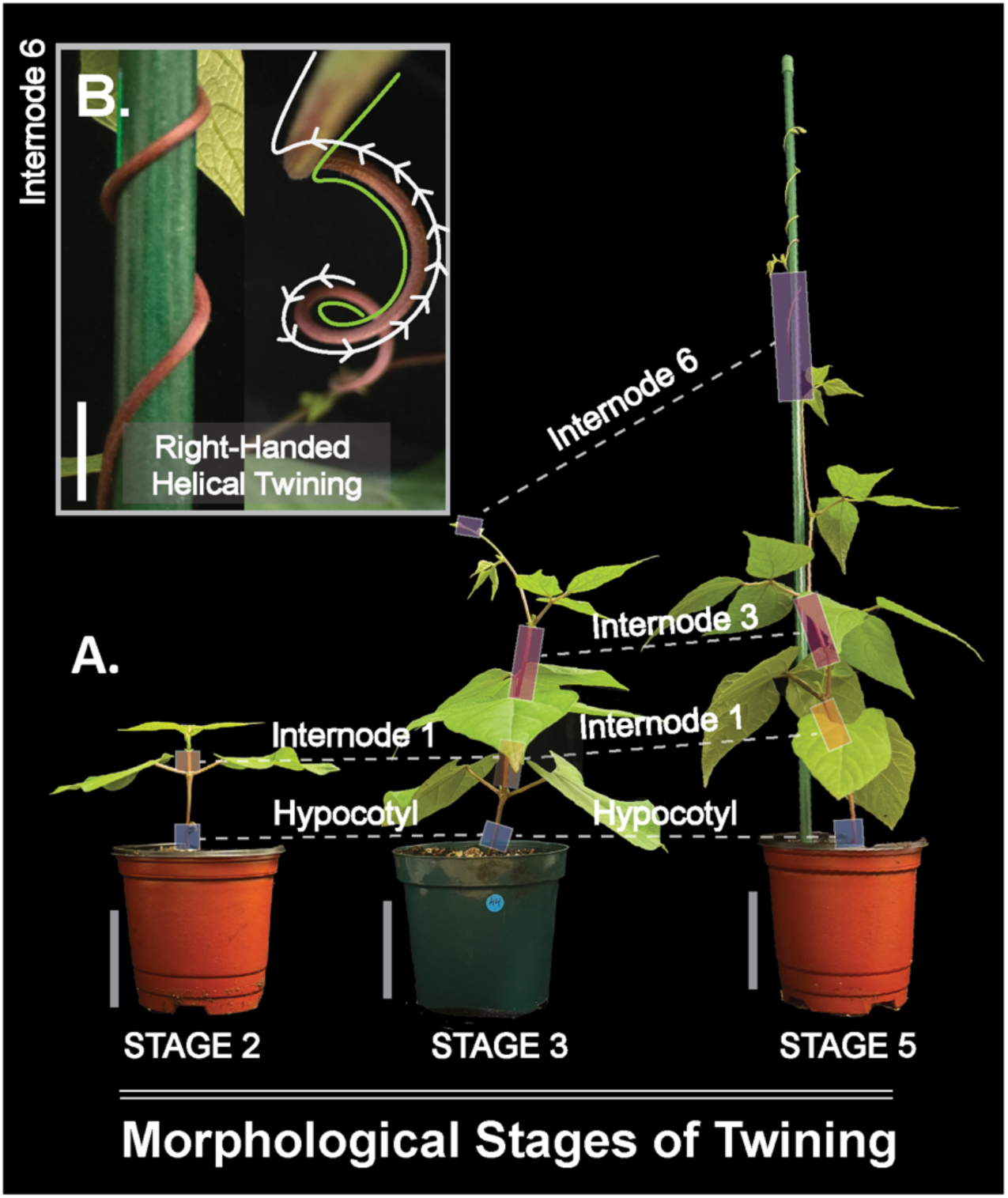
Morphological stages of twining in common bean. (**A**) Morphological stages of twining in common bean sensu Onyenedum et al. (2025). Each sampled segment, the hypocotyl, internode 1, internode 3, and internode 6, was highlighted with a box throughout developmental stage. (**B**) Close-up of a right handed twining at internode 6, stage 5. Scale bars, A = 5 cm, B = 6.25 mm.

**Fig. 2:**
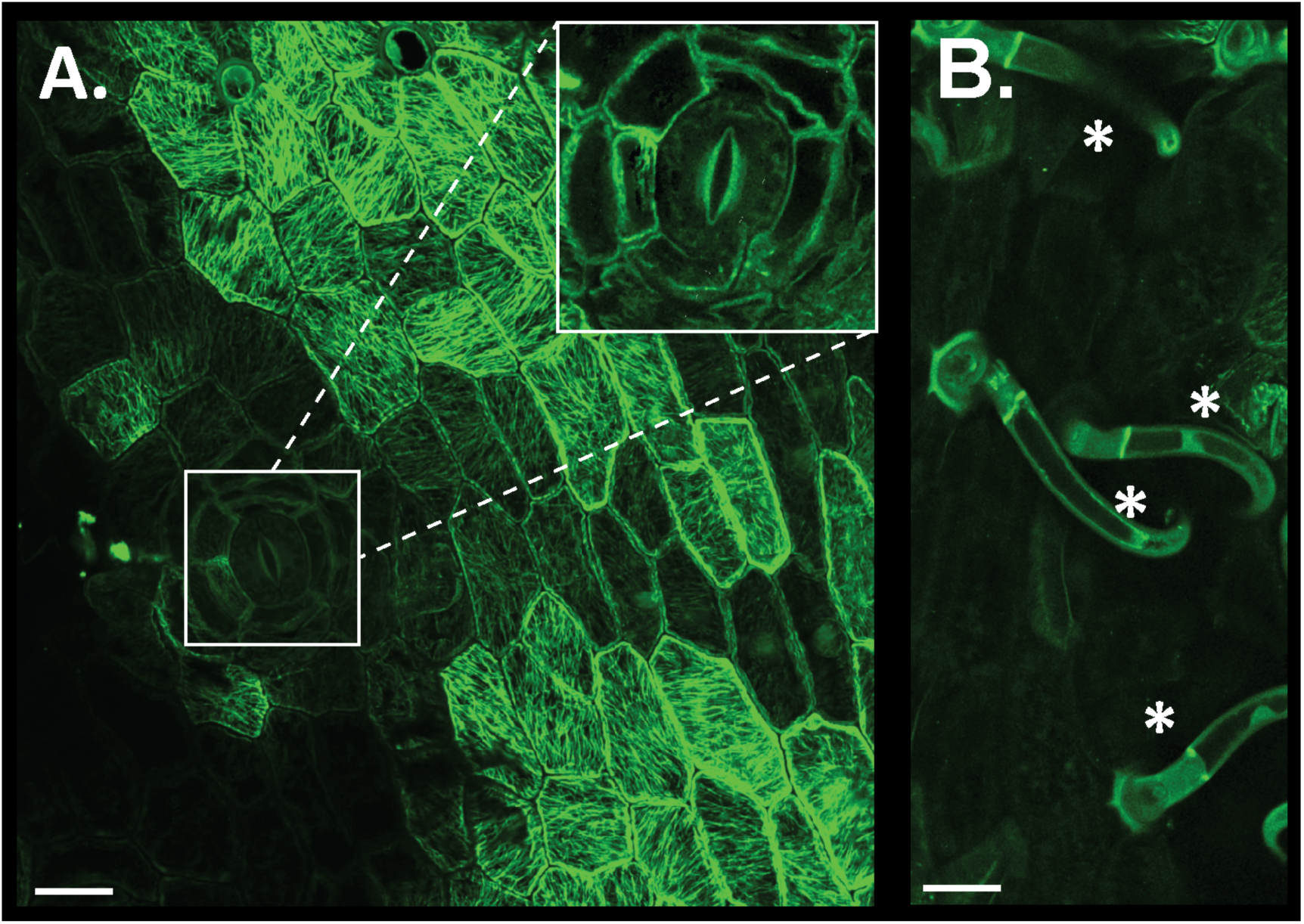
Cortical microtubule (CMTs) arrays along the periclinal face of epidermal cells in common bean. Epidermal markers, such as (**A**) stomata and (**B**) trichomes (*) were used as reference points to locate the periclinal face of sampled epidermal peels. Immunostaining was performed using an anti-α-tubulin antibody excited by a 488nm laser. Scale bars = 25 μ*m*.

**Fig. 3:**
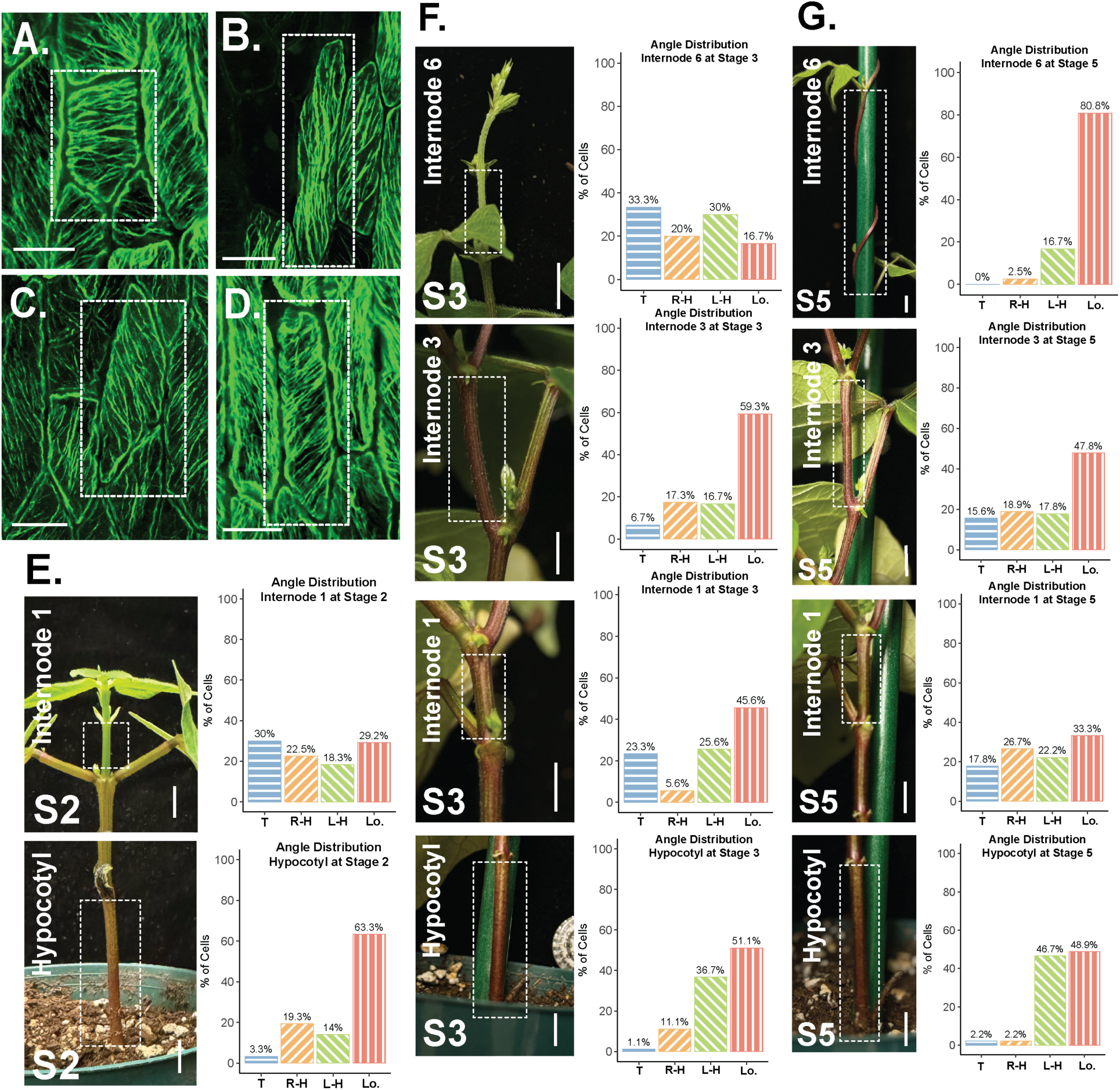
Cortical microtubule angle bins, and their distribution in the hypocotyl, internode 1, internode 3, and internode 6 through three morphological stages of common bean. (**A–D**) CMT arrays binned based on the net orientation of the outermost cell wall: (**A**) transverse CMTs, (**B**) longitudinal CMTs, (**C**) left-skewed CMTs, and (**D**) right-skewed CMTs. (**E–F**) Reference stem segment photos provide detail into the organ configuration of straight and twisted regions of a developing common bean vine through morphological stages—(**E**) stage 2 (S2), (**F**) stage 3 (S3), (**G)** stage 5 (S5). Barplots described the makeup of CMT patterning based on the percentage of orientation groups across morphological stages: transverse (0 – 22.5° and 157.5 – 180°), right-skewed (22.5 – 67.5°), longitudinal (67.5 – 112.5°), and left-skewed (112.5 – 157.5°). Scale bars A–D = 25 μ*m*, and E–G = 1cm. T = Transverse orientation; R-H = right-handed skew; L-H = left-handed skew; Lo. = longitudinal orientation.

**Table 1.** Microtubule Orientation Chi-square and corresponding residuals. A total of 30 cells were measured for each biological replicate (90 cells = 3 replicates, 150 cells = 5 replicates).

| Stage | Tissue | Chi-square residuals |  |  |  | Dominant Category / Significance |
| --- | --- | --- | --- | --- | --- | --- |
|  |  | Transverse | Right-skewed | Longitudinal | Left-skewed |  |
| S2 | Hypocotyl | -6.13 | -1.6 | 10.84 | -3.11 | Longitudinal (highly significant) |
| S2 | Internode 1 | 1.26 | -0.63 | 1.05 | -1.69 | No significant deviation |
| S3 | Hypocotyl | -5.23 | -3.04 | 5.72 | 2.56 | Longitudinal (significant) |
| S3 | Internode 1 | -0.37 | -4.26 | 4.5 | 0.12 | Longitudinal (significant) |
| S3 | Internode 3 | -5.19 | -2.17 | 9.71 | -2.36 | Longitudinal (highly significant) |
| S3 | Internode 6 | 1.83 | -1.1 | -1.83 | 1.1 | No significant deviation |
| S5 | Hypocotyl | -4.99 | -4.99 | 5.23 | 4.75 | Longitudinal, Left-skewed (significant) |
| S5 | Internode 1 | -1.58 | 0.37 | 1.83 | -0.61 | No significant deviation |
| S5 | Internode 3 | -2.07 | -1.34 | 4.99 | -1.58 | Longitudinal (significant) |
| S5 | Internode 6 | -6.32 | -5.69 | 14.12 | -2.11 | Longitudinal (highly significant) |

\* Residuals > |±1.96| indicate significant deviation at $p < 0.05$ . Residuals > |±2.58| indicate significant deviation at $p < 0.01$ .

### CMT are mostly mixed and longitudinal in expanding and fully elongated internodes, respectively

#### Stage 2 CMT orientation of stationary basal stem segments

In stage 2, young plants are erect and short and comprised of a stationary aboveground hypocotyl, epicotyl, and internode 1 (Fig. 1A, “Stage 2”). At this stage, we compared the CMT orientations in the fully elongated hypocotyl (mean ± SD = 2.67 cm ± 0.47 cm, n = 7), to the newly emergent internode 1 (mean ± SD = 1.34 cm ± 0.33 cm, n = 7). In the hypocotyl (Fig. 3E, “Hypocotyl”), CMT orientations were primarily longitudinal (63.33%), with significant overrepresentation confirmed by Chi-square analysis (Table 1). By contrast, there was no dominant angle category for CMT within the short developing internode 1 (Fig. 3E, “Internode 1”; Table 1).

#### Stage 3 CMT orientation of dynamic and intermediary stem segments

In stage 3, plants underwent regular circumnutation and were comprised of ∼ seven internodes (Fig. 1A, “Stage 3”). The hypocotyl, epicotyl, internode 1, and internode 2 were stationary, while internodes 4–7 underwent regular circumnutation, characterized by oscillating movement of the shoot. Internode 3 was an intermediary stem segment between the stationary base and dynamic apical internodes. By this stage, internode 1 (mean ± SD = 2.18 cm ± 0.46 cm, n = 9) and internode 3 (mean ± SD = 3.91 cm ± 1.09 cm, n = 9) had reached their full length, while internode 6 (mean ± SD = 1.93 cm ± 1.11 cm, n = 9) was still elongating (Fig. S1).

Like the previous stage, the straight hypocotyl’s dominant CMT orientation remained longitudinal (51.11%) (Fig. 3F, “Hypocotyl”; Table 1). In fully elongated internode 1, the CMT orientations shifted to a dominant longitudinal (45.6%) CMT orientation (Fig. 3F, “Internode 1”; Table 1). The fully elongated internode 3 had a dominant longitudinal (59.33%) CMT orientation (Fig. 3F, “Internode 3”; Table 1). Lastly, the newly emerged internode 6 had mixed CMT orientations, with no dominant angle (Fig. 3F, “Internode 6”; Table 1).

#### Stage 5 CMT orientation of the matured twined stem

In stage 5, basal segments, such as the hypocotyl and internode 1, remained straight and stationary. Internode 3, the intermediary segment, transitioned from straight and dynamic to straight and stationary (Onyenedum et al., 2025). Internode 6, the most apical region, transitioned from short and circumnutating to elongated and twined with the assistance of a stake (mean ± SD = 10.53 cm ± 3.19 cm, n = 14) (Fig. S1).

Compared to the previous two stages, the straight hypocotyl’s dominant CMT orientation shifted away from being longitudinally dominant to overrepresentation of both longitudinal (48.89%) and left-skewed (46.67%) (Fig. 3G, “Hypocotyl”; Table 1). Internode 1 returned to a mixed grouping, with no dominant CMT orientation (Fig. 3G, “Internode 1”; Table 1). Internode 3 CMT orientations remained significantly overrepresented in the longitudinal category (47.8%) (Fig. 3G, “Internode 3”; Table 1). Finally, within the twined internode 6, CMT orientations was significantly overrepresented in the longitudinal category (80.83%) (Fig. 3G, “Internode 6”; Table 1).

Broadly, internodes that had not yet completed elongation (internode 1 at stage 2 and internode 6 at stage 3) displayed a mix of CMT orientations, with the largest presence of transverse CMT grouping compared to other stages (Fig. 3E, 3F). In contrast, segments that had just completed elongation, including the hypocotyl (Stage 2), internode 1 (Stage 3), internode 3 (Stage 3), and internode 6 (Stage 5), all displayed a dominant longitudinal CMT orientation (Fig. 3E – 3G; Table 1). Post-elongation stages of hypocotyl (Stage 3, Stage 5) and internode 1 (Stage 5) revealed a subtle transition from longitudinal patterning to an increased proportion of skewed orientations (left– and/or right-skewed). Additionally, it can be noted when tracking skewed orientation groups at stage 5, internodes 1 and 3 possessed relatively equal proportions of left– and right-skewed CMT orientations, whereas the hypocotyl and internode 6 were biased towards left-skewed orientations.

### Undulated stem morphology corresponds to the distribution of oblong and rounded cells

While sectioning each stem segment throughout stages 2, 3, and 5, we observed variations in cross-sectional morphology of the hypocotyl, internode 1, internode 3, and internode 6 (Fig. 4A). While the hypocotyl emerged circular and remained circular, each succeeding internode emerged slightly undulated, containing protrusions associated with cortical tissue surrounding the vascular bundles and pericyclic fibers. The protrusions, referred to as “lobes,” were separated by shallow grooves termed “furrows” that run along the main axis of the stem (Fig. 4B, C). This lobular morphology begins mildly in internode 1, whereas the hypocotyl remains entirely circular (Fig. 4A, 4C). Given the realization that the stem circumference was not homogenous, we sought to understand if there is a relationship between the configuration of the internode (straight vs. coiled), the contrast between lobes/furrows, cell shape, and CMT orientation. Towards this aim, we sampled twined plants in stage 5 and characterized cell morphologies (cell area, circularity, and geodesic diameter) and CMT angles at internode 1, internode 3, and internode 6, separating the “lobes” from the “furrows.” Hypocotyls were excluded given the lack of undulation.

**Fig. 4:**
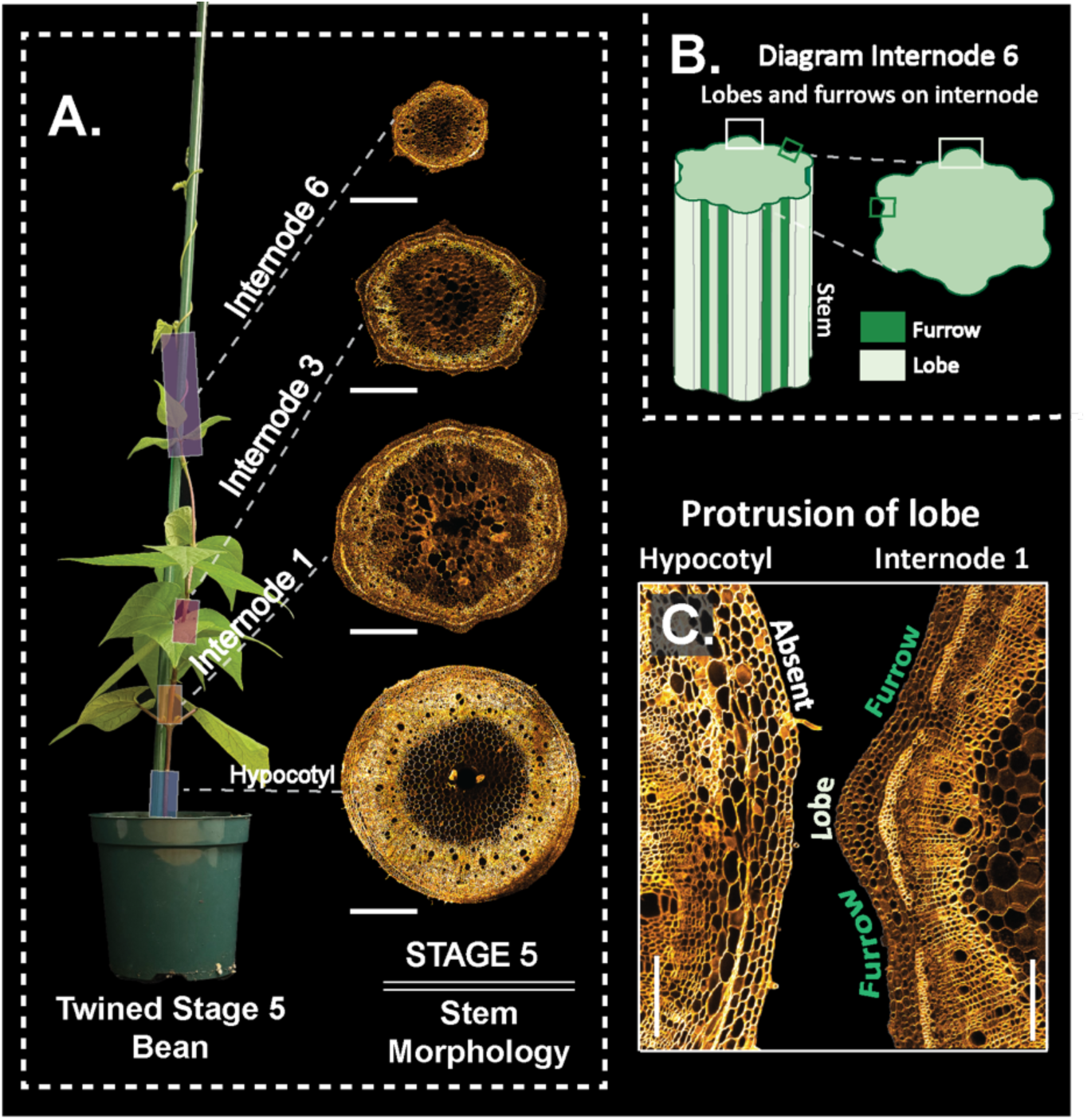
Characterizing stem morphology at stage 5 of common bean. **(A)** Cross-sections of stem segments to illustrate the circular outline of the hypocotyl, and progressively undulated outline from internode 1 to internode 3, to internode 6. **(B)** Illustration of the longitudinal and cross-sectional distribution of lobes and furrows at internode 6. **(C)** A cross-sectional comparison of a hypocotyl and internode 1 illustrates a lack of protrusions found around the periphery of the hypocotyl. Note the furrows and lobes in internode 1. Scale bars, A = 500μ*m*, C = 125 μ*m*.

In internode 1, area of epidermal cells in the furrows and lobes were not statistically distinct (p-value = 0.68). However, cells in the furrow were more round (significantly higher circularity (p-value < 2e-16) than those in the lobes, and had a smaller geodesic diameter (p-value < 2e-16), a measure of length of the maximal axis (Fig. 5A, 5B). In internode 3, cell areas remained similar between the lobes and furrows (p-value = 0.56), while furrows maintained a higher circularity (p-value < 2e-16) and significantly smaller geodesic diameter than the lobes (p-value = 5.6e-11; Fig. 5C, 5D). In the twined internode 6, both cell areas (p-value = 0.068) and geodesic diameter (p-value = 0.068) were statistically indistinguishable, but the cells in the furrows continued to exhibit higher circularity values than those in the lobes (p-value = 9.9e-05; Fig. 5E, 5F), although this distinction was less pronounced than in internodes 1 and 3.

**Fig. 5:**
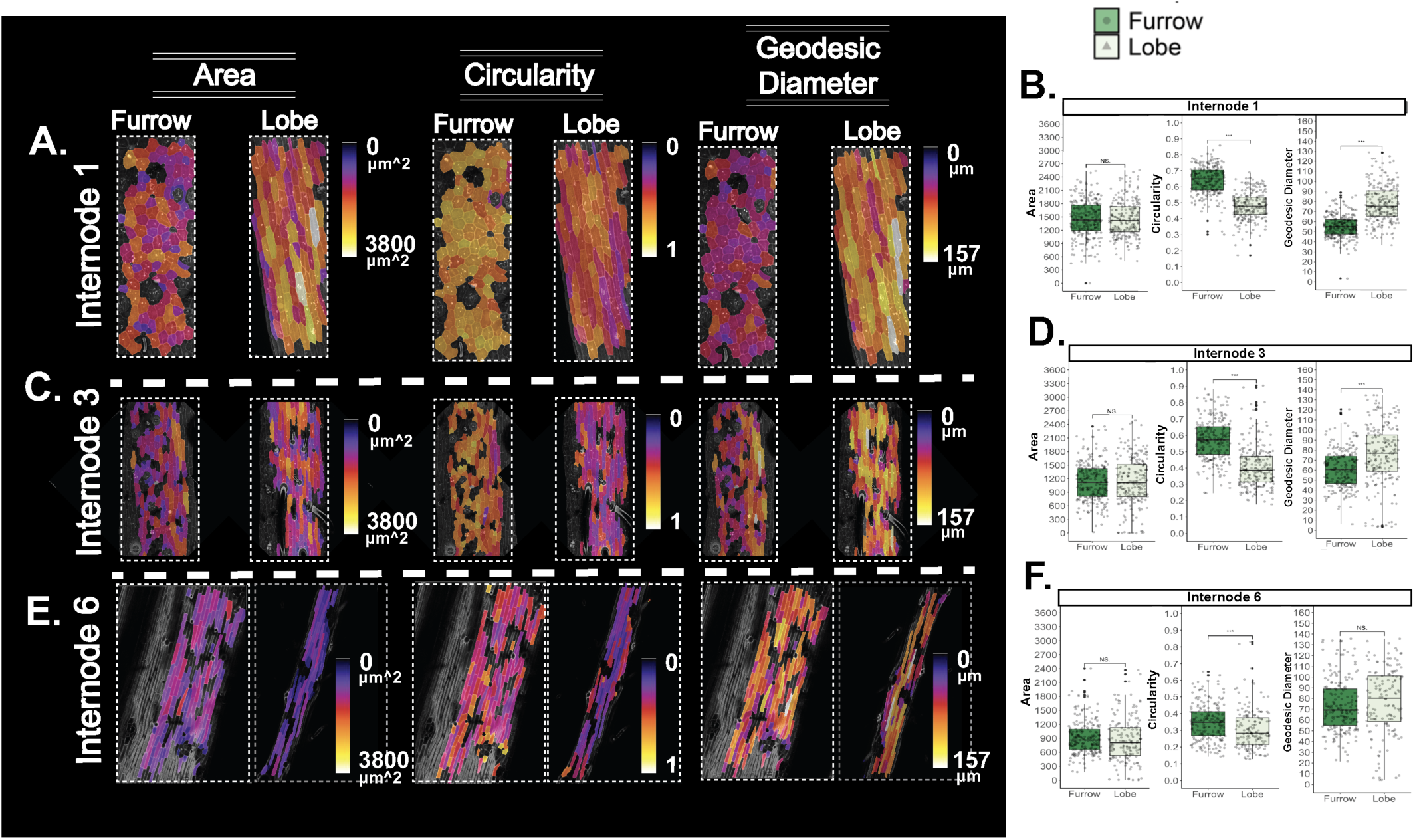
Cell morphologies across furrows and lobes in twined common bean (stage 5). Epidermal peels of **(A)** internode 1, **(C)** internode 3, and **(E)** internode 6 were segmented based on their lobed or furrowed positions. Individual segmented cells were measured based on their cell area (0 – 3800 μ*m*”), circularity (scale 0 to 1) describing the ‘roundness’ of the cell, and geodesic diameter (0 – 157 μ*m*), quantifying the length of the longest (maximal) axis of the cell accounting for cell convexity. Wilcoxon statistical tests were used to test significance values between lobes and furrows on (**B)** internode 1 (**D**), internode 3, and (**F**) internode 6.

Taken together, the cell area is the same across both furrows and lobes in all three internodes examined, thus the morphological distinction between those regions in internodes 1 and 3 comes down to a distinction in circularity and geodesic diameter. Interestingly, those distinctions became less pronounced or eliminated in internode 6, where cells were more rectangular, appearing “stretched” (Fig. 6A).

**Figure 6:**
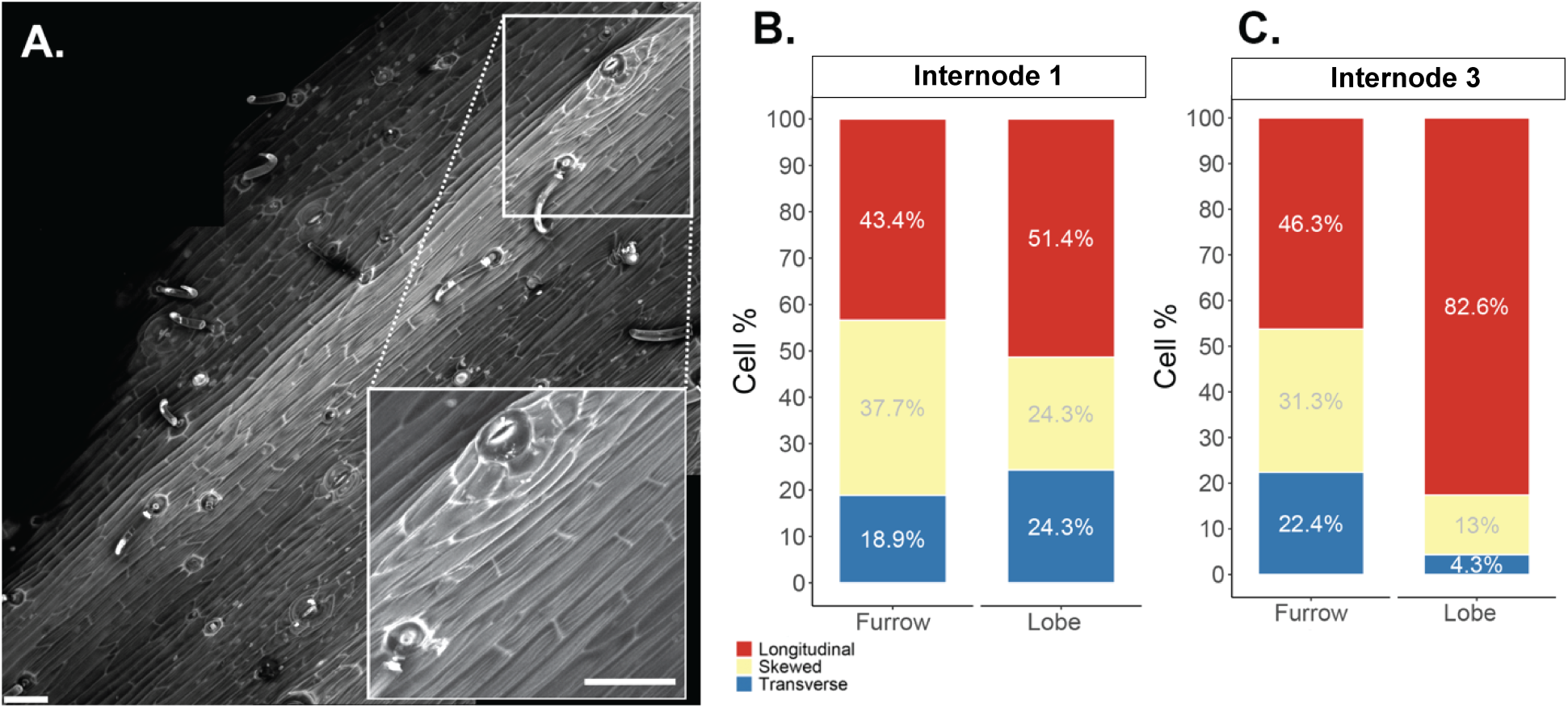
Cell morphologies and cortical microtubule angles in common bean. (**A**) Highlight of rectangular cell morphologies found along a twined internode 6 in stage 5. The vertical striations along the cell wall are presumably caused by the stretching of the organ while climbing. Image of sample stained with 10µg/mL propidium iodide (PI). Scale bar = 50 μ*m*. **(B, C)** Stacked bar plots illustrate the representation of each cortical microtubule orientation bin in stage 5 plants. **(B)** Internode 1 furrows versus lobes. **(C)** Internode 3 furrows versus lobes.

To evaluate the association between CMT orientation and cell morphogenesis, epidermal cells were grouped based on their position within the furrows or lobes of internodes 1 and 3 in twined stage 5 plants. These internodes were selected due to the significant differences in circularity and geodesic diameter observed between the lobes and furrows (Fig. 5). By analyzing these regions separately, we found that within internode 1, the furrows and lobes both displayed a mixed patterning of CMT angles with no contrast in overall distribution (p-value = 0.73, Wilcoxon test) besides a slight overrepresentation of longitudinal CMTs in the lobes according to chi-square residual testing (Fig. 6B; Table 2). Internode 3 furrows and lobes differed in overall distribution (p-value = 0.0062), but both had an overrepresentation of longitudinal orientation at 46.3% and 82.6%, respectively (Fig. 6C; Table 2).

**Table 2.**
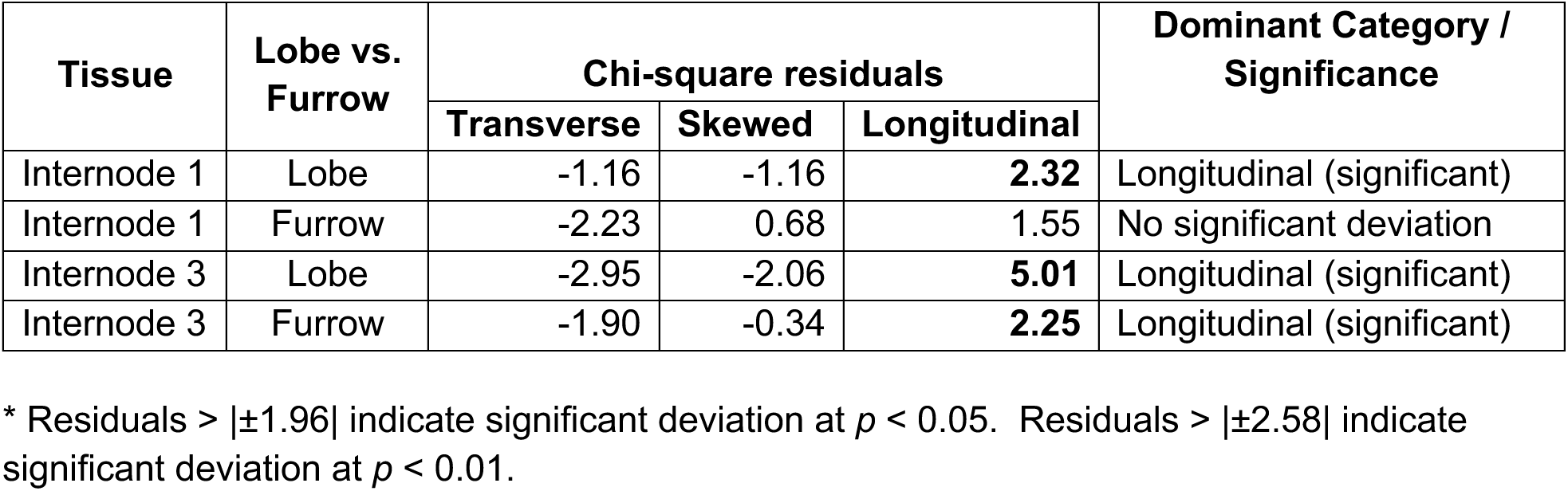
Stage 5 Microtubule Orientation Chi-square and corresponding residuals for lobes and furrows. Cell Measurements from biological replicates in Table 1 were regrouped based on their positioning in either ‘lobed’ or ‘furrowed’ regions of the stem.

## Discussion

This study draws inspiration from the prevailing hypothesis in the literature that the helical growth of *Arabidopsis thaliana* twisted mutants is shaped by the cytoskeleton. A major conclusion derived from this literature is that cortical microtubules are implicated in skewed helical growth through microtubule-associated proteins (e.g., *SPIRAL1*, *tortifolia*/*SPIRAL2*), α– and β-tubulin mutations (e.g., *TUA*, *TUB*), and microtubule-disrupting drug sensitivity (e.g., propyzamide, *PSH1*) (Nakajima et al., 2006; Ishida et al., 2007; Naoi and Hashimoto, 2004). By sampling plants expressing a twisted phenotype, a consensus has emerged that the skewed orientation of microtubules is often perpendicular to the direction of cell and overall organ growth (Smyth, 2016; Buschmann and Borchers, 2020). The first and, to our knowledge, only work examining microtubule orientation in a naturally twining species was conducted in *Ipomoea nil* with a preliminary dataset strikingly similar to twisted *Arabidopsis* mutants (Sousa-Baena et al., 2021). These works helped to form our initial hypothesis that the right-handed growth of common bean vines likely traced back to a predominance of left-skewed microtubules in the epidermis––the main tissue layer responsible for organ formation (Kutschera et al., 2007; Wada, 2012). We approached this study by sampling not only twined plants at maturity, but also re-sampling the four stem segments (Hypocotyl, Internodes 1, 3, and 6) throughout plant maturation from a seedling (Stage 2) to early circumnutation (Stage 3) to a coiled adult (Stage 5). Therefore, our results not only inform us about the association between microtubule orientation and coiled internodes but also about how microtubule patterning changes as an internode elongates, undergoes regular circumnutation, and eventually coils.

### Does skewed microtubule orientation predict and reflect organ directionality?

To our surprise, we did not find a dominance of skewed microtubules at any stage, thus refuting our initial hypothesis. Instead, we found that fully elongated internodes, both straight and coiled, exhibited predominantly longitudinal microtubules. Meanwhile, still elongating internodes displayed a mixture of arrangements, with the highest frequency of transversal orientation occurrence (Fig. 3E, “Internode 1”; Fig. 3G, “Internode 6”). This observation provides support for earlier findings that cortical microtubules and cellulose microfibrils often align transversely to the growth axis (Green, 1962), a phenomenon also noted in ten-day-old *Ipomoea nil* hypocotyls (Sousa-Baena et al., 2021). However, in the twined internodes of common bean, we found a left-skewed over right-skewed bias in the hypocotyl and internode 6, albeit this left-skewed bias was always second place to the dominant longitudinal pattern (Fig. 3G, “Hypocotyl” & “Internode 6”). Interestingly, both stem segments demonstrate the capacity for twisting—just as observed in the hypocotyl during shoot emergence from the soil (Fig. S2), and across internode 6 which couples twisted and twined forms (Fig. 1B). This preference aligns with existing knowledge of directional cell expansion (Baskin, 2005), suggesting a link between left-skewed microtubule orientation and right-handed cell-files. In contrast, straight stem segments, internodes 1 and 3, were relatively at equilibrium between right– and left-skewed microtubules, with the only exception being internode 1 (Stage 3) when it is undergoing circumnutation (Fig. 3E–3G). If skewed microtubule orientation indeed reflects organ chirality in twining vines, our findings suggest it emerges later in development, after elongation. Nonetheless, given the dominance of longitudinal microtubule patterning found, we will focus on alternative hypotheses explaining this pattern below.

### Longitudinal microtubules may reflect tension across stretched twined internodes

So why are cortical microtubules aligned longitudinally in twined internodes? As discussed above, twined internode 6 poses a high percentage of left-skewed over right-skewed microtubules, but this is only second to the dominant pattern of longitudinally arranged microtubules (>80%) (Fig. 3G, “Internode 6”). A possible explanation is that cortical microtubules dynamically rearrange in response to external stimuli, such as mechanical damage (Sampathkumar et al., 2014), nutrient acquisition (Sheng et al., 2024), and/or tension and compression patterns across plant tissue (Hejnowicz et al., 2000; Hamant et al., 2008). Significant tensile and compressive forces applied to plant tissue can cause microtubules to realign parallel to the direction of maximal force (Sahaf et al., 2016; Robinson and Kuhlmeier, 2018; Zhao et al., 2020; Li et al., 2023).

As a twining plant ascends a support structure, tensile stresses naturally occur. A circumnutating stem must supply ample grip to secure contact with a support and counteract gravitational forces, consequently stretching the stem along the length of the support (Fig. 6A) (Silk and Hubbard, 1991; Silk and Holbrook, 2005; Isnard et al., 2009). According to the main stages of tension identified by Isnard et al. (2009) in stipule-assisted climbing, *Discorea bulbifera* can experience up to 150 to 200nM in squeezing forces when coiled around a pole and distant from the shoot apex. Considering the similarities of *D. bulbifera* to twining common bean, we predict that twined internode 6 likely undergoes similar squeezing forces. In *Arabidopsis*, tensile and compressive forces as low as 1.5 to 2mN were sufficient to induce microtubule realignment from transverse to longitudinal in the hypocotyl (Robinson and Kuhlmeier, 2018). Therefore, if common bean experienced comparable tensile forces induced by climbing a host, these stresses could drive substantial microtubule rearrangement.

### Longitudinal microtubules may reflect secondary growth of the stationary internodes

Longitudinal patterning in basal, straight stem segments like the hypocotyl, internode 1, and internode 3, cannot be readily explained by twining-related tension. Instead, we propose that the expansion of inner tissue by secondary growth drives this microtubule arrangement. As the common bean stem thickens post-elongation, vascular expansion from secondary growth, exerts pressure on outer tissues, particularly the epidermis. The epidermal growth theory (“tensile skin theory”) highlights the bidirectional stresses exchange between inner and outer tissues (Kutchshera, 1989; Hejnowicz and Sievers, 1995). Supporting this theory, studies on sunflower hypocotyls demonstrated an increased inner tissue growth of 15% due to the release of external epidermal constraints (Hejnowicz et al., 2000; Kutschera et al., 2007). These findings emphasize the epidermis’ role in shaping organs and managing axial growth forces exerted by internal tissues. However, in radial expansion, the thickening of vascular tissue is capable of generating circumferential strain on the stem and serving as a growth directional cue for the organ, similar to commonly studied axial growth (Baskin and Jensen, 2013). Thus, post-elongation longitudinal microtubule arrangement likely facilitates epidermal vertical expansion, enabling the tissue to counteract internal radial pressure from secondary growth while preserving the structural integrity of the organ.

### Distribution of oblong and rounded cell morphologies across the stem periphery

While examining common bean morphology, we observed a transition from a circular basal stem to a slightly undulating, noncircular stem in distal internodes (Fig. 4A). Protrusions along the stem periphery correlated with a thicker cortex and positioning of pericyclic gelatinous fibers, which generate localized areas of tensile stress and reinforce stem posture (Melloerowicz, et al., 2008; Mellerowicz and Gorshkova, 2012; Onyenedum et al., 2025). The transition between stem morphologies may serve distinct purposes in posture maintenance: the circular basal hypocotyl has a prominent role during self-supporting growth, while the more distal internodes develop protruding “lobes” and recessed “furrows” (Fig. 4A) that may assist with early flexibility required in twining. Their shape is similar to other climbing plants with non-circular stems, such as *Bauhinia* sp., *Passiflora coccinea,* and *Paullina* sp. (Bhambie, 1972; Rowe & Speck, 2005; Isnard & Silk, 2009; Chery et al., 2020). These implications emphasize the functionality of “lobes” and “furrows” concerning twining, and prompted us to assess their composition at the cellular level. Along the stem perimeter, we also found distinct cell morphologies between the lobes and furrows. Lobes contained rectangular-shaped cells, while furrows had round-shaped cells, the latter becoming relatively more rectangular and stretched in apically twined internodes (Fig. 5A, 5C, 5E; Fig. 6A). These differences suggest variations of isotropic and anisotropic cell growth between the two regions—however, microtubule organization patterns found suggest an added layer of complexity to cell formation. During post-elongation, stage 5, internode 1 lacked a clear distinction in microtubule orientation between the lobe and furrow (Fig. 5B); meanwhile, internode 3’s lobe demonstrated a notably higher affinity for longitudinal arrangement than the furrow with 82.6% and 46.3%, respectively (Fig. 6C). These observations suggests that common bean stems can compartmentalize cell growth to create a structure that is initially self-supporting and then well-adapted for climbing. However, we were unable to pinpoint microtubule arrangement as the definitive origin of lobes and furrows variability; the complexity lies in the dynamic nature of microtubule arrangement that shifts amidst plant development, stress, damage, or other key factors.

### Free-standing twisting organs vs helical-twining along a support

Although *Arabidopsis thaliana* mutants have laid the groundwork for investigating microtubule-driven twisting, key differences exist between “twisting” versus “twining”. A useful analogy is a shoestring fixed at one end: in twisting, the free end is twirled, creating a drill-bit shape. In contrast, twining occurs when the shoestring stretches its axis, wrapping around a support, forming a spring-like coil. *Arabidopsis* twisted mutants exhibit skewed directional growth at the cellular and organ levels, evidenced by twisted epidermal cell files (Thitamadee et al., 2002, Ishida et al., 2007) and individual cells (Verger et al., 2019). In contrast, twining plants like common bean must elongate and slightly twist as a mechanical necessity when twining (Darwin, 1875; Isnard et al., 2009). This twisting is less pronounced than in *Arabidopsis* mutants, where millimeter-sized organs exhibit drastic cell file skewing (Furutani et al., 2000; Thitamadee et al., 2002; Nakajima et al., 2004). Instead of extreme cell file twisting, twined epidermal cells appear more longitudinally stretched (Fig. 6A), likely due to organ-level stress imposed during climbing. Moreover, twining plants do not rely solely on twisting to maintain their helical structure—they require constant support contact for early posture adherence via thigmotropism (Huberman and Jaffe, 1986; Onyenedum et al., 2025). For instance, once circumnutating internodes establish unilateral contact with a stake, emerging internodes use the support as a guide, simultaneously twining and elongating into a coiled position. Taken together, the clear dominance of skewed microtubules in *Arabidopsis* and the lack therefore in common bean, may reflect two different modes of helical growth: cells grow in a twisted manner to form twisted organs in *Arabidopsis*, versus the internode must both elongate and grow in a helical fashion in contact with a support in common bean.

By examining a fully twined stem, we investigated microtubule orientation in the context of the complete twining phenotype rather than isolated instances of twisting. Furthermore, the added nuances of the twining plant form increased the complexity of our question considering the sensitivity and dynamics of microtubules. While the exact relationship between climbing-induced tissue stress and microtubule rearrangement remains unclear, our findings lay the groundwork for exploring how tensile stress influences longitudinal MT arrangements in twining mechanics.

## Supplemental

**Supplemental Figure 1:**
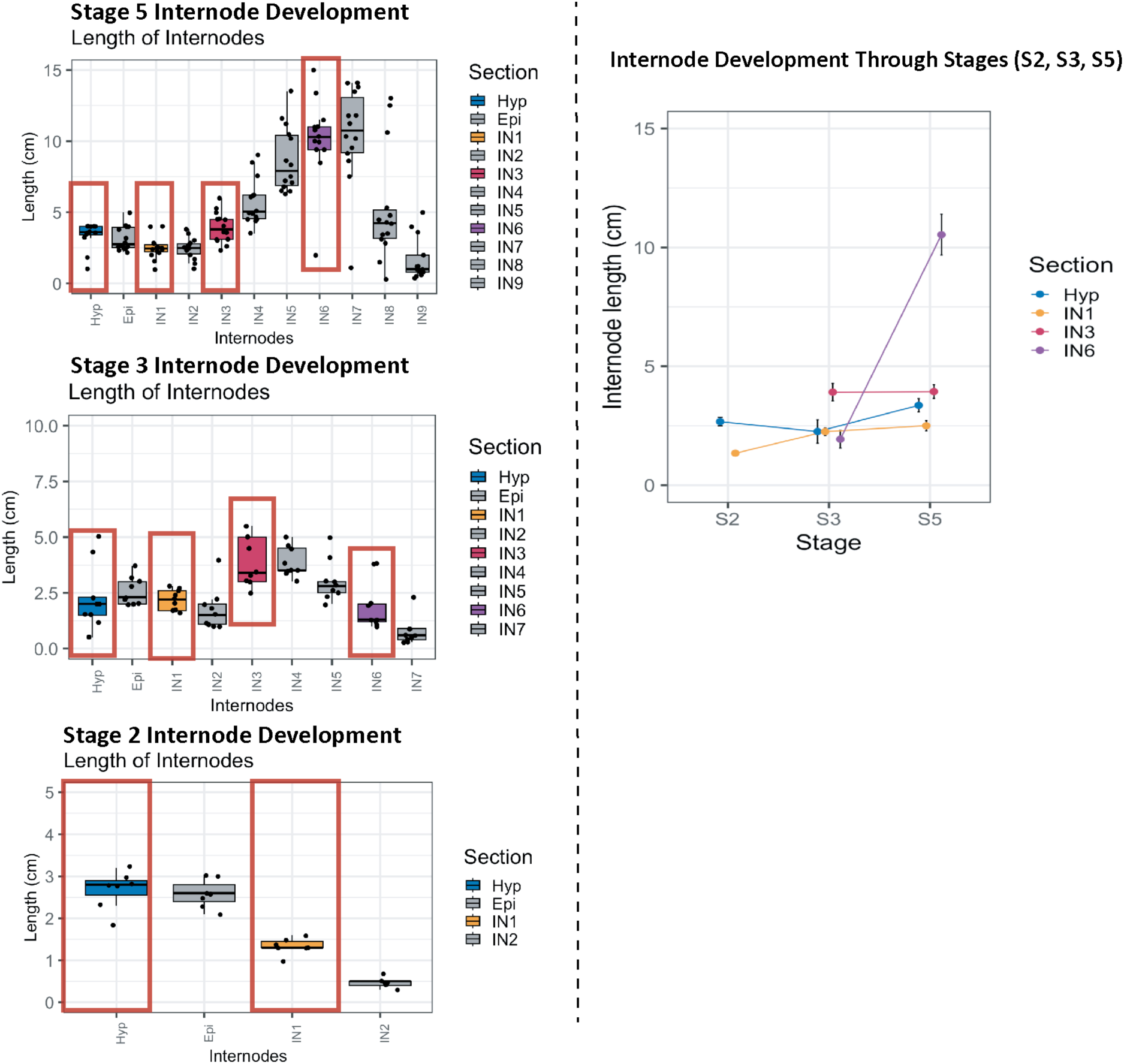
Internode development throughout stages 2, 3, and 5. At stage 2, the hypocotyl has completed elongation, while internode 1 is still elongating. By stage 3, the hypocotyl, internode 1, and internode 3 have all completed elongation, but internode 6 has just initiated. By stage 5, internode 6 has elongated to an average of > 10 cm and fully reached a twined state. Red boxes denote the regions of interest in microtubule analysis.

**Supplemental Figure 2:**
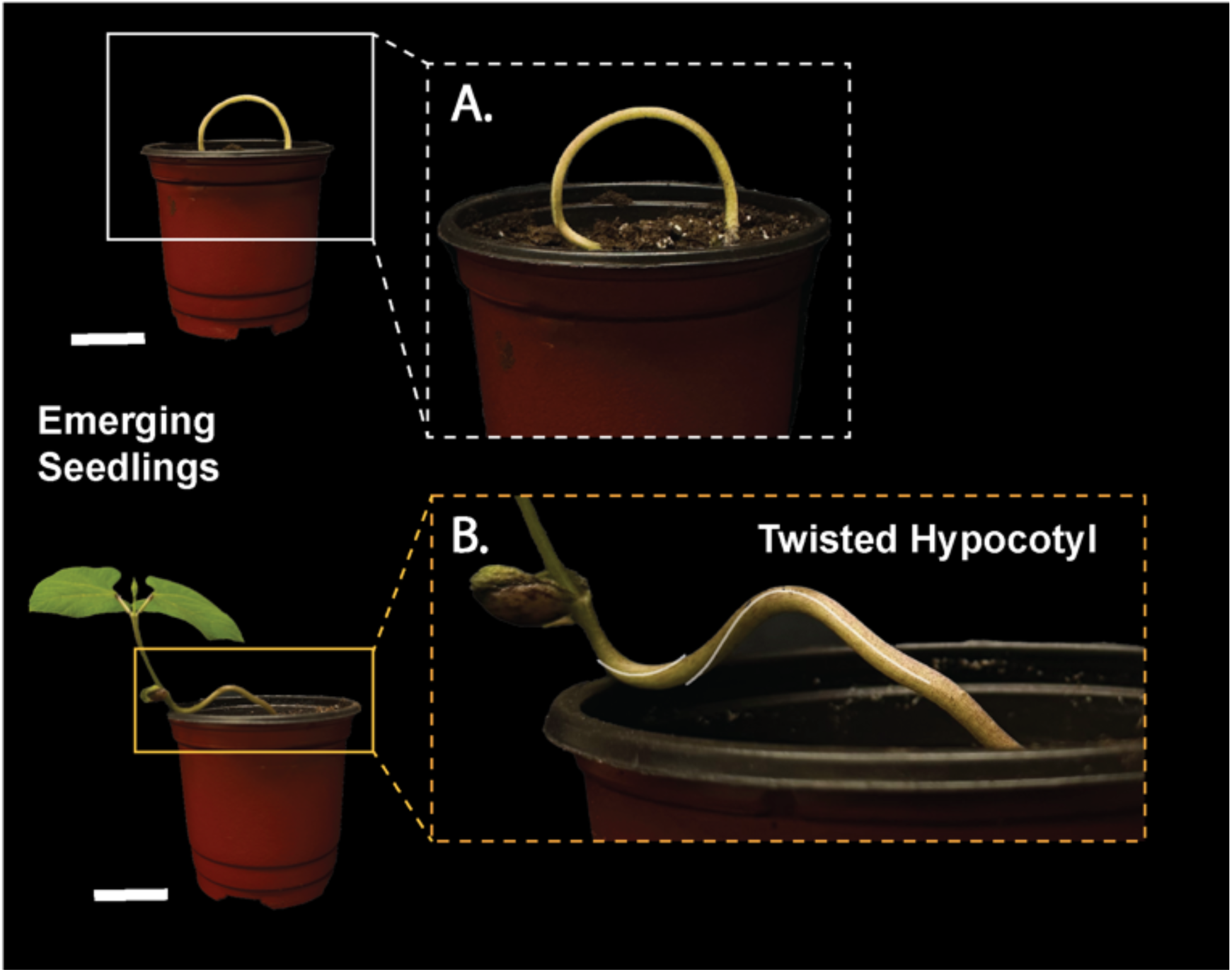
The hypocotyl of common bean seedlings has the capacity to be twisting when guiding or repositioning the shoot apex upward toward the light source. This resembles the documented self-twisting nature of *Arabidopsis* twisted mutants.

**Supplemental Figure 3:**
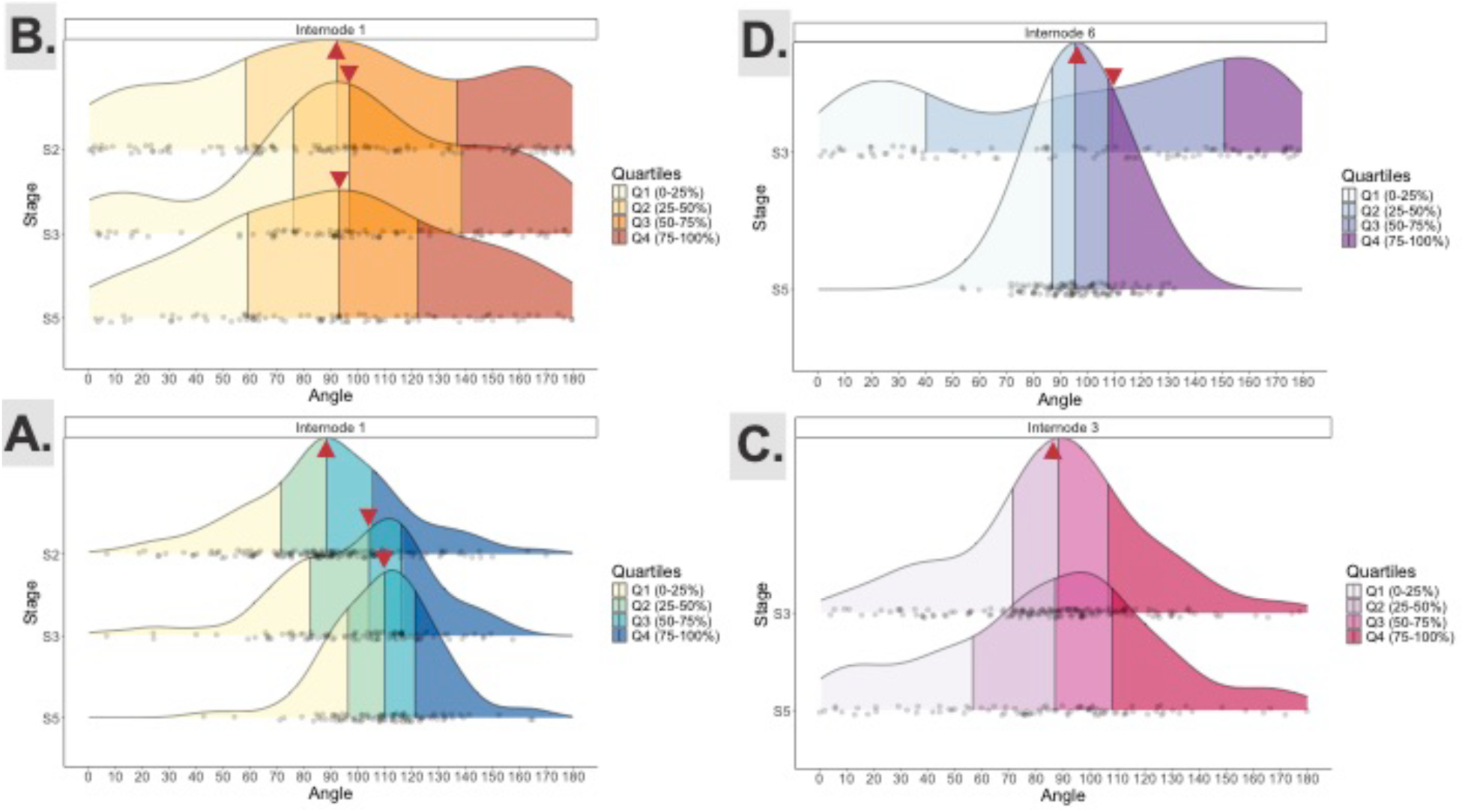
Ridgeline relative density and distribution plots detail the broad distribution of CMT orientations measured at. (A) the hypocotyl, (B) internode 1, (C) internode 3, (D) internode 6 of stages 2, 3, and 5, while providing insight into the quartile ranges (Q1, Q2, Q3, Q4) and standard deviations of the dataset. Red triangles point to the median.

## Acknowledgements

A special thank you to all Onyenedum lab members—Lena Hunt, Israel L. Cunha-Neto, Zachary Kozma, Hannah Ratcliff, Yifan Wang, Annabelle Wang, and Leo Semana—for their unconditional support and advice with experiments, revisions, and, of course, plant care.

## Author Contributions

J.G.O, M.S.S.B, and A.A.A conceptualization; M.S.S.B and A.A.A methodology; A.A.A investigation; J.G.O and A.A.A formal analysis; J.G.O resources; A.A.A data curation; J.G.O and A.A.A scripts; J.G.O and A.A.A writing – original draft; A.A.A, M.S.S.B, and J.G.O review & editing; J.G.O and A.A.A visualization; J.G.O supervision; J.G.O funding acquisition.

